# Computational Lead Optimization on BACE1: Relative Binding Free Energy Perturbation as the Terminal Refinement Layer

**DOI:** 10.64898/2026.07.07.737131

**Authors:** Kristoffer Alejo, Christopher Korban, Christian Chung

## Abstract

Structure-based drug discovery is known to apply computational methods in a tiered hierarchy, with each layer narrowing the candidate set and refining the binding picture before committing to the next, more expensive step. We present a four-tiered computational benchmarking study evaluating five engines against a panel of 36 compounds targeting *β*-secretase 1 (BACE1), a validated Alzheimer’s disease target with extensive co-crystal ground truth. This study evaluates Flexible Docking and Boltz2 Cofolding as the primary tier, followed by Ensemble Docking, and then Protein-Ligand MD with MM/PBSA and MM/GBSA post-processing. This is then concluded with Relative Binding Free Energy Perturbation (RevFEP) as the terminal refinement layer. Each method was benchmarked against the experimental binding free energies derived from the co-crystal structures spanning −7.85 to −11.35 kcal/mol. Our findings revealed that Flexible Docking reproduced the co-crystal binding mode for 35 of 36 ligands (97.2% within 2.0 Å RMSD) but did not rank potency at this resolution. Boltz2 CoFolding provided an orthogonal structural cross check with a receptor backbone RMSD of 0.293 Å against the experimental co-crystal structure. Ensemble Docking identified the optimal receptor conformation for downstream FEP setup. MD with MM/GBSA decomposition identified van der Waals complementarity as the primary potency driver (Pearson *r* = +0.855, *R*^2^ = 0.732 on a 10-compound subset). RevFEP delivered the highest affinity correlation of any method (Pearson *r* = +0.662, *R*^2^ = 0.438, Spearman *ρ* = +0.624, mean absolute error 1.02 kcal/mol across all 36 ligands), resolving potency differences within a narrow 3.5 kcal/mol congeneric window that no other engine could discriminate. We characterize what each engine contributes independently and where RevFEP delivers signals no other engine achieves.

## 1 Introduction

Alzheimer’s Disease is one of the most widely known neurodegenerative disorders characterized by severe cognitive impairment symptoms disruptive to daily life Tenchov et al. (2024). It is caused in large part by the extracellular accumulation of amyloid plaques and the intracellular formation of neurofibrillary tau tangles Abdulkhaliq et al. (2026), which collectively drive neuroinflammation, synaptic loss, and severe disruptions in cholinergic, glutamatergic, and GABAergic neurotransmissions Abdulkhaliq et al. (2026) Sun et al. (2009). The amyloid precursor protein (APP) is characterized in its role in the brain synapse preserving dendritic spine morphology in addition to regulating the neurotransmission of both glutamate and GABA Kruse et al. (2025). In the Alzheimer’s disease state, the processing of APP shifts towards the amyloidogenic pathway, where instead of *α*-secretase separating the protein within the Amyloid-*β* sequence, B-secretase (BACE1) cleaves APP at the *β*-site, producing a membrane-bound fragment subsequently cleaved by *γ*-secretase to release Amyloid-*β* peptides into the extracellular space, where they accumulate and aggregate into the amyloid plaques characteristic of Alzheimer’s disease Zhao et al. (2020). BACE1 represents the primary therapeutic intervention point, where inhibition upstream prevents Amyloid-*β* production before aggregation can occur Koelsch (2017).

The BACE1 structure consists of an active site where the hydrogen bonding interaction at the ASP32 and ASP228 residues facilitate the binding between the protein and molecule Ghosh and Osswald (2014). Access to the binding site is controlled by the flap region, which is defined structurally as a *β*-hairpin loop. This flap region controls whether the protein assumes an open conformation, where access to the active site makes ligand binding favorable, or a closed conformation, where the flap restricts access, rendering the ligand unable to bind to the active site Shimizu et al. (2008). The BACE1 active site is also composed of two large hydrophobic subpockets in the S1 and S3 regions Ghosh and Osswald (2014), where pharmacophoric design prioritizes facilitating binding in this specific region Patil et al. (2010). The primary limitation of BACE1 lead optimization is designing a molecule that is small enough and lipophilic enough to cross the Blood-Brain Barrier (BBB), while also maintaining enough contacts in a large pocket to achieve a high binding affinity Imran et al..

In any drug discovery campaign, given an identified target, the main computational bottleneck becomes lead optimization. Thousands of compounds can be proposed for synthesis; however, synthesis and assay resources are finite. As a result, the main question that needs to be addressed is, given a large compound library, how can we decide which ligands to prioritize for synthesis and assay testing? Previous studies have been conducted on the BACE1 target as part of a lead optimization campaign. In Wang et al., Relative Binding Free Energy (RBFE) has been established as a viable tool for industrial lead optimization, validating Schrodinger’s FEP+ engine on a set of 36 compounds designed to target BACE1 Wang et al. (2015). This set of 36 compounds all consisted of an amidine-containing core scaffold, with experimental binding affinities spanning a tight range of 3.5 kcal/mol from -7.85 to -11.35 kcal/mol Wang et al. (2015). In this study, FEP+ achieved a mean unsigned error of 0.84 kcal/mol and a correlation of r = 0.78 Wang et al. (2015). Though FEP is known to be incredibly accurate, it is largely limited by the intensity of compute resources, limiting the scalability of this particular tool Ross et al. (2023).

In this study, we propose a multi-stage computational benchmarking study, utilizing different engines from pose generation to measuring high-resolution potency differences. The engines utilized here range in terms of accuracy and computational costs. The five engines evaluated in this lead optimization campaign are: Flexible Docking, Boltz2 CoFolding, Ensemble Docking, Molecular Dynamics (MD) with MMPBSA/GBSA postprocessing, and Relative Binding Free Energy Perturbation (RBFE). Flexible Docking and CoFolding operate in parallel as the primary layer, with the goal of pose generation using model based methodology and physics based methodology respectively. Both outputs can then be hedged against one another to validate the binding geometry Nguyen et al. (2020) Passaro et al. (2025) Liu et al. (2026). Ensemble docking follows with the purpose of identifying the optimal protein conformation for ligand binding Totrov and Abagyan (2008) Huang and Zou (2007). Protein-Ligand MD with MMPBSA/GBSA is then conducted to characterize the thermodynamic drivers of binding Genheden and Ryde (2015). RBFE serves as the terminal refinement layer delivering high resolution potency rank ordering at the sub-kcal/mol level Muegge and Hu (2023). In this paper, we benchmark five computational engines of varying accuracy and computational cost, each evaluated independently against experimental binding affinities. We aim to demonstrate how each method contributes non-redundant information towards sub-kcal/mol potency discrimination with the end goal of synthesis prioritization in lead optimization in mind. This protocol aims to minimize computational costs compared to heavier pure FEP+ calculations as well. Each of these engines would be utilized beginning with initial target ID and identification of hits all the way to down-selection and lead optimization during an iterative Design-Make-Test-analyze (DMTA) Cycle.

**Figure 1:**
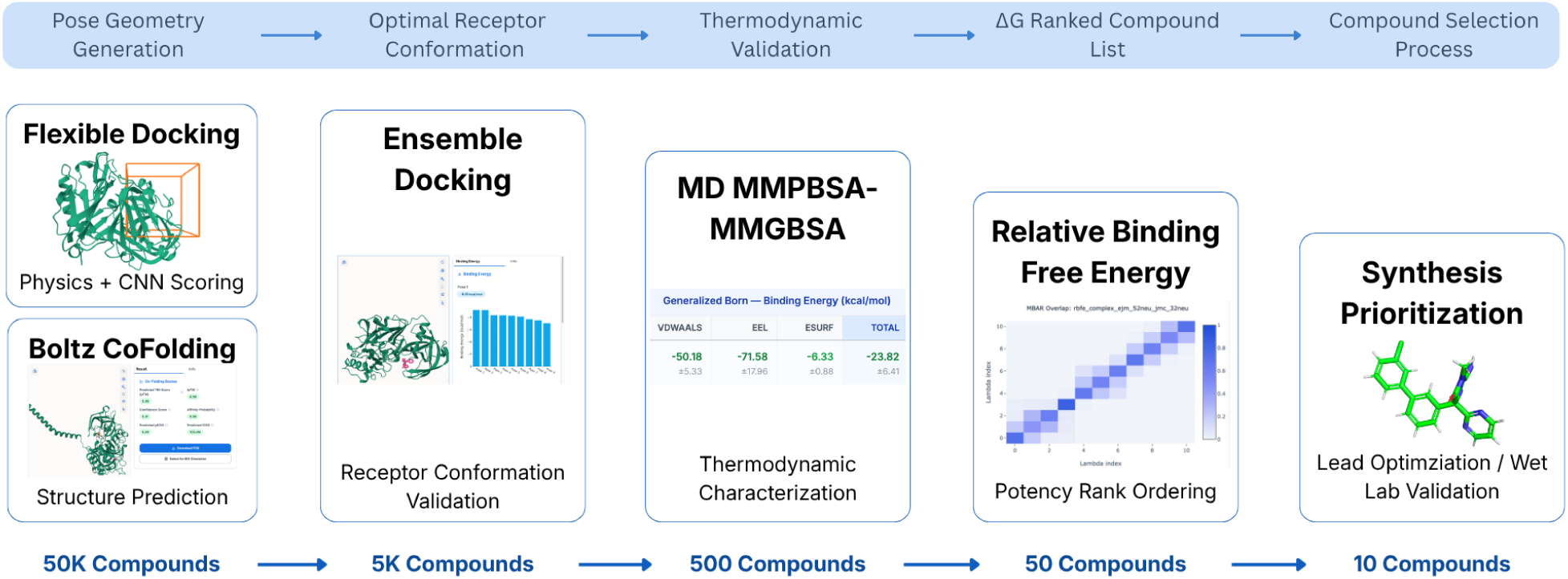
Overview of the five computational engines applied to the BACE1 congeneric series. Left to right, the engines becomes increasingly more computationally rigorous and accurate, with each engine evaluated on experimental binding free energy.

## 2 Methods

The compound set evaluated in this study is the Schrödinger BACE1 benchmark dataset referenced and utilized in Wang et al. (2015). As mentioned above in the introduction section, there are a total of 36 ligands in this set, all containing an amidine-containing core scaffold. Experimental ΔG in this set ranged from -7.85 to -11.35 kcal/mol. The dataset is composed of 36 total co-crystal structures of each ligand bound to the BACE1 receptor. Structural data consisted of a single BACE1 receptor PDB file (PDB: 4DJW) and an SDF file containing all 36 ligand structures with experimental ΔG in kcal/mol stored in the (r exp dg) property field. Co-crystal 3D poses are available for all 36 ligands, enabling the use of the dataset for RMSD validation. Further preprocessing prior to downstream analysis was performed. The protein PDB file was first preprocessed using PyMOL Rosignoli and Paiardini (2022). The protein preprocessing protocol contains the following steps: removal of water molecules from the crystal structure, removal of co-crystallized ligand for apo structure preparation when needed, removal of heteroatoms not relevant to the binding site, addition of missing hydrogens, inspection and repair of missing residues or broken chains if applicable. The cleaned structure is then exported as a PDB file for downstream use. Ligand preprocessing included the segmentation of the single SDF file containing all 36 ligands into 36 separate SDF files, each containing a single ligand. Lastly, experimental ΔG was then converted to pIC50 with the following equation:

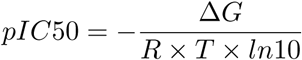

Wang et al. (2015)

Flexible Docking was then performed using the cleaned 4DJW PDB protein input as well as the 36 individual SDF files from the segmentation step. Docking grid parameters were set to a size of 30Å x 30Å x 30Å. For defining the docking grid center coordinates, a manual inspection of each of the 36 co-crystal structures was performed, ensuring that all compounds were bound to the receptor in the same pocket. The centroid of each ligand’s 3D coordinates was calculated, then averaged across all 36 ligands to obtain the single docking grid center coordinates. The engine was run with an exhaustiveness of 32, representing 4x the default sampling length, through more random restarts and search iterations. For each ligand evaluated, 9 poses were returned with a total of 324 poses generated. The scoring function for binding was evaluated using both physics scoring and CNN rescoring Meli et al. (2022). The top pose per ligand was selected based on the CNN score. RMSD validation was also performed comparing the top pose for each ligand against the experimental co-crystal structure, where RMSD ≤ 2.0 Å was the baseline for determining a successful pose reproduction. MCS-based atom mapping via RDKit was utilized to compare the atomic positions between structures Fooshee et al. (2013). The primary outputs to be used in downstream analysis are the CNN affinity score, the physics docking score, and the RMSD per ligand.

Boltz CoFolding was then performed in parallel to flexible docking. The protein sequence for BACE1 was obtained from Uniprot using the PDB ID: 4DJW, where the isoform P56817-1 was identified as the canonical structure and utilized as the protein sequence input. The panel of 36 SMILES strings was processed with RDKit, converting the 3D ligand structures from the SDF file into SMILES strings for each ligand. The purpose of Boltz CoFolding is to predict the 3D protein-ligand complex and binding affinity from both sequence and SMILES alone, without a pre-defined binding pocket. A single 3D co-crystal structure, a confidence score per prediction, and predicted affinity outputs in the form of pIC50 metric are then outputted. Analysis on the prediction accuracy of Boltz CoFolding was performed by evaluating predicted IC50 values against experimental IC50, originally converted from experimental ΔG measurements. In addition, structural validation analysis was performed on the Boltz2 predicted complex for CAT-13a, as CAT-13a served as the anchor ligand in the downstream FEP perturbation network as well as sat at the mid-range of the experimental potency distribution making it a representative compound of the full set. The Boltz2 predicted complex and the co-crystal structure were aligned along Chain A using Pymol, where protein backbone RMSD was computed via the align function and ligand pose RMSD was computed via the pair-fit function on heavy atoms only. It is also worth noting that no model fine-tuning was performed on the BACE1 dataset; instead only the base Boltz2 model was used for evaluation.

Ensemble Docking was performed downstream of both flexible docking and Boltz CoFolding. Before Ensemble Docking, Protein Water Molecular Dynamics was performed on the apo BACE1 structure. The engine generated a trajectory of 100 ns with an integration timestep of 0.002 ps. The AMBER99SB-ILDN force field Lindorff-Larsen et al. (2010) was applied in addition to the TIP3P Water Model Mark and Nilsson (2001). Solution utilized NaCl with a concentration of 0.15 M, where the system was heated to 300K for equilibration. The generated MD trajectory was then preprocessed before ensemble docking, segmenting the trajectory into 10 equidistant snapshots, converting the 100ns trajectory into 10ns intervals. Ensemble docking was performed with the following inputs: the apo BACE1 pdb file and the BACE1 ligands sdf file. For each of the 10 snapshots, a 30Å x 30Å x 30Å was generated as the docking grid. Grid center for each snapshot was defined manually by referencing structural landmarks of the BACE1 binding pocket, specifically secondary structure elements including alpha helices and *β* sheets flanking the active site Dislich and Lichtenthaler (2012), as each snapshot is fundamentally different from the apo protein structure.The engine was run with an exhaustiveness of 8, reduced from flexible docking due to multiplied compute costs. Each of the 36 ligands was then docked against each of the 10 receptor snapshots. 9 poses were generated for each prediction, resulting in a total of 3240 poses. The primary metric for evaluation is the best CNN score across all 10 snapshots per ligand, with the mean CNN score as the secondary metric, with the rationale that the best score captures the most favorable binding conformation sampled. The aggregated CNN affinity score and aggregate physics score were then utilized in downstream analysis.

Protein-Ligand Molecular Dynamics with MMPBSA/GBSA postprocessing was performed downstream of ensemble docking. Before execution, a subset of 10 compounds from the set of 36 BACE1 compounds was equidistantly sampled based on experimental ΔG to ensure a representative sample of the entire dataset. This subset size was determined by the substantial computational cost of running 100 ns MD per ligand, where scaling to the full 36-compound set would require 3.6 times the compute at approximately 3.6 *µ*s of aggregate simulation time, which was prohibitive within the scope of this study. Molecular dynamics trajectories were then generated utilizing the experimental co-crystal structures for each of the 10 compounds sampled. Although docked poses were generated, the co-crystal structures were used as starting geometries for MD simulations to maintain proper thermodynamic characterization, independent of any uncertainty generated from the previous pose generation stages Wang et al. (2015). This practice is consistent with the standard practice for MMPBSA and MMGBSA benchmarking. The trajectories were generated using the AMBER99SB-ILDN forcefield Lindorff-Larsen et al. (2010), and the TIP3P water model Mark and Nilsson (2001). NaCl solution with a concentration of 0.15 M was utilized, where the system was heated to 300K for equilibration. A trajectory of 100 ns was generated for each of the 10 co-crystal structures, where frames were saved every 100 ps, yielding 1000 frames per trajectory. Each frame was extracted from the production trajectory, and MMPBSA and MMGBSA were then computed for each ligand. Binding free energy was then outputted and decomposed for each component (e.g., van der Waals, electrostatic, solvation contributions, etc.). Downstream analysis was then performed comparing predicted ΔG against experimental ΔG for each component calculated. To assess for potential confounding of MMGBSA VDW correlation with ligand size, we computed the pearson r between molecular weight and experimental ΔG for each of the 10 ligands in our set. Benjamini-Hochberg false discovery rate correction was applied across the five following energy components to correct for multiple comparisons: MMGBSA VDW, MMGBSA EGB, MMGBSA EEL, MMGBSA TOTAL, MMPBSA TOTAL.

Relative Binding Free Energy Perturbation (RBFE) was then performed on all 36 ligands in the Schrödinger BACE set using Revilico’s FEP engine. Before the execution of any RBFE protocol, a perturbation network was constructed using all 36 ligands from the set. The network was produced using the minimum spanning tree network topology alongside the Kartograff atom mapper Ries et al. (2024). In total, there were 70 transformations produced. RBFE was then performed with 11 total *λ*-windows, with an MD trajectory of 5 ns being produced at each *λ*-window for each leg (an MD trajectory produced for the ligand-bound state and a trajectory for the ligand in solvent) across all 70 transformations. MD was calculated using AM1-BCC Jakalian et al. (2002) as the charge method, OpenFF 2.0 (Sage) Boothroyd et al. (2023) as the small molecular force field, Amber ff14SB Maier et al. (2015) as the protein force field, and TIP3P Mark and Nilsson (2001) as the water model. Multistate Bennett Acceptance Ratio (MBAR) Ding (2024) was then utilized for free energy estimation across *λ* windows. Binding Free Energy was then calculated as the difference in free energy between the complex leg and the ligand leg, and can be calculated with the following equation:

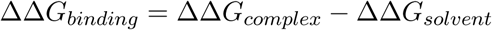

Wang et al. (2015)

The output relative ΔΔG values were then converted to absolute predicted ΔG using maximum likelihood estimation, as MLE minimizes the squared deviation across the full perturbation network. Per-transformation uncertainty estimates were also reported. Per-transformation uncertainty estimates were then computed via MBAR and reported alongside predicted ΔΔG values.

## 3 Results

RMSD validation of generated poses from flexible docking was performed for all 36 ligands against the co-crystal structures, where successful pose reconstruction was defined as an RMSD < 2.0 Å. 35 of 36 ligands were reconstructed below the 2.0 Å threshold, with CAT-24 identified as the sole outlier with an RMSD of 2.36 Å (Figure 2A). The RMSD distribution across the full set assumes a tight, approximately normal or slight right skew, centered around 1.0 Å, with no bimodal behavior (Figure 2A). The Schrödinger BACE set was assessed with a mean of 1.16 Å, a median of 1.13 Å, a minimum of 0.5 Å, and a maximum of 2.36 Å (Table 1). Affinity correlation analysis was then performed using Pearson r and Spearman *ρ* to evaluate flexible docking affinity outputs against the experimental pIC50 values. Four metrics were compared against experimental pIC50 values: Best CNN Affinity (pK), CNN Affinity at Best Pose Score (pK), Best Physics Score (kcal/mol), and Affinity Gap (kcal/mol). Affinity gap is computed using the difference between the best physics-based affinity and the mean physics-based affinity. Both CNN-based methods showed little rank correlation with Best CNN Affinity evaluated with r = -0.155, *ρ* = -0.174, and CNN Affinity at Best Pose Score evaluated with r = 0.039, *ρ* = 0.025 (Figure 2B). The physics-based metrics showed a stronger but inverse correlation with Best Affinity evaluated with r = -0.454, *ρ* = -0.434, and Affinity Gap with r = -0.306, *ρ* = -0.350.

**Figure 2:**
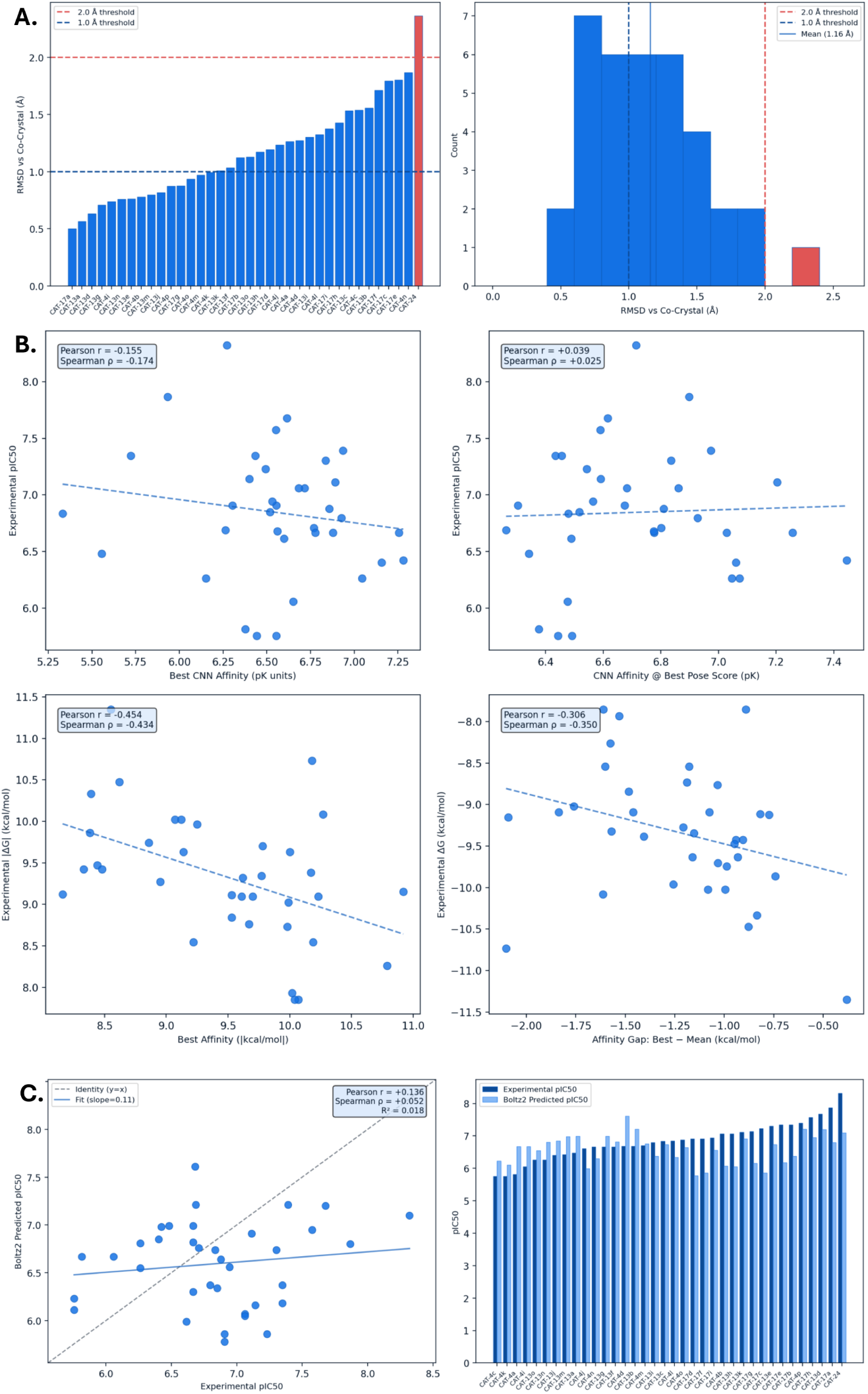
Flexible Docking and Boltz CoFolding. **A)** Flexible docking pose reproduction against co-crystal structure (N=36). Left: per-ligand RMSD sorted ascending, with 35 of 36 ligands below the 2.0 Angstrom threshold. Right: RMSD distribution, mean 1.16 Angstroms. CAT-24 (red) identified as only ligand above threshold. **B)** Flexible docking score versus experimental affinity (N=36). No docking metric shows meaningful rank correlation with potency; they physics score is inverted and CNN affinity is near zero. Top Left: Best CNN Affinity (pK) vs Experimental pIC50. Top Right: CNN Affinity at Best Pose Score (pk) vs. Experimental pIC50. Bottom Left: Best Affinity (kcal/mol) vs Experimental |DeltaG| (kcal/mol). Bottom Right: Affinity Gap calculated using Best – Mean (kcal/mol) vs Experimental DeltaG. **C)** Boltz CoFolding predicted vs. Experimental pIC50 (BACE1, N = 36). Left: Predicted vs Experimental pIC50 scatter plot fit with regression line.. Right: Per-Ligand Predicted pIC50 bar graph sorted from weakest to strongest binder.

**Table 1:** Boltz2 CoFolding structural validation for CAT-13a against experimental co-crystal structure (PDB: 4DJW).

| Analysis | RMSD (Å) | Threshold (Å) |
| --- | --- | --- |
| Protein Backbone | 0.293 | < 1.0 |
| Ligand Pose | 4.082 | < 2.0 |

All 36 ligands from the Schrödinger BACE series were evaluated with Boltz2 CoFolding. Specifically, predicted pIC50 was evaluated against experimental pIC50 using Pearson r, Spearman *ρ*, and *R*^2^. Results show experimental pIC50 ranges from 5.76-8.32 compared to predicted pIC50 ranges from 5.78-7.61. Experimental pIC50 vs predicted pIC50 has little to no correlation with r = 0.136, *ρ* = 0.052, and *R*^2^ = 0.018 (Figure 2C). Figure 2C displays the discrepancy between experimental pIC50 values and predicted pIC50 values at the ligand-specific resolution. Notably, CAT-24 was identified as the largest underprediction in the series, with a difference of 1.22 pIC50 units. Structural validation performed on CAT-13a resulted in a protein backbone RMSD of 0.293 Å and a ligand pose RMSD of 4.082 Å (Table 1). Confidence scores ranged from 0.88 to 0.91 and affinity probability from 0.92 to 0.98 across all 36 ligands, with both scores assuming a tight and high scoring range, indicative of no discrimination across the series.

All 36 ligands were docked across 10 receptor snapshots of a Protein-Water MD trajectory for BACE1. Receptor 3 was identified as the geometrically optimal conformation with a mean CNN Affinity of 6.31 pk and CNN Pose Score of 0.76 vs the dataset mean of 0.35 (Figure 3B). Receptor 3 also obtained a strain pass rate of 100 percent, where strain pass is defined as a Best Intramol score ≤ 0 kcal/mol. Receptor 2 was identified as the least productive conformation with a CNN Affinity of 4.38 pK and CNN Pose Score of 0.23 (Figure 3B). Receptor 10 obtained a strain failure rate of 16.7 percent, the highest rate across all snapshots. Receptor 8 was the only snapshot with a marginally positive Spearman *ρ* of 0.04 (Figure 3B). Affinity correlation did not improve over flexible docking at the ensemble level. Three aggregation metrics were evaluated against experimental pIC50. Ensemble Best CNN Affinity was evaluated with r = -0.229, *ρ* = -0.262, Ensemble Mean CNN Affinity with r = -0.147, *ρ* = -0.205, and Ensemble Best Physics Affinity with r = -0.419, *ρ* = -0.408. No aggregation metric achieved meaningful affinity correlation. The physics score reached statistical significance but with an inverted rank direction (*ρ* = -0.408).

**Figure 3:**
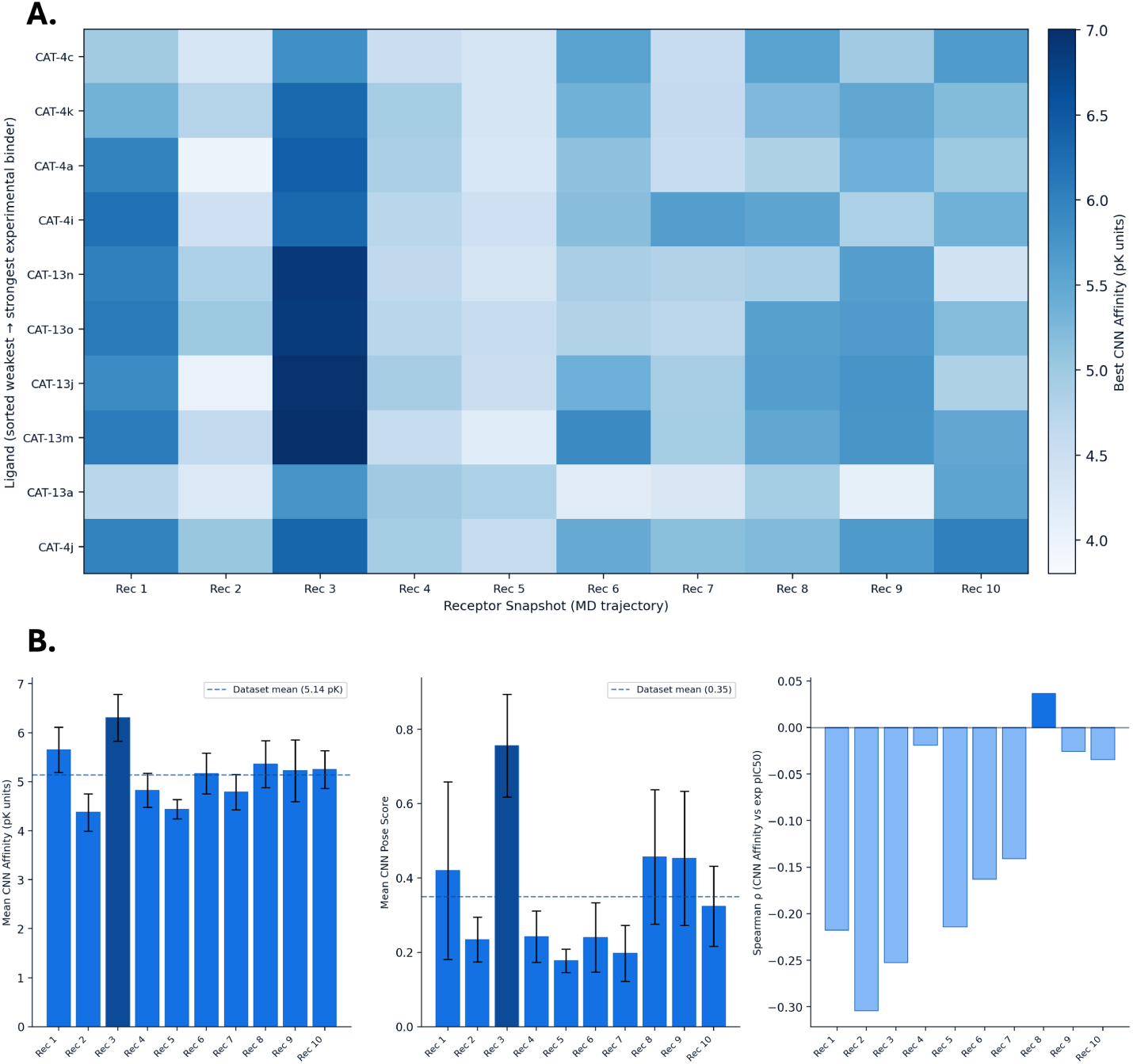
Ensemble Docking. **A)** Ensemble docking CNN affinity heatmap truncated to 10 ligands across 10 receptor screenshots. Heatmap with full 36 ligand panel across 10 receptor screenshots provided in supplementary figure section below. Receptor 3 (column 3) shows the strongest predicted binding across the series. **B)** Receptor snapshot quality summary. Left: Mean CNN Affinity for each receptor. Higher CNN Affinity = stronger predicted binding. Middle: Mean CNN Pose Score for reach Receptor. Higher CNN pose score = more realistic pose geometry. Right: Rank Spearman Correlation vs Experimental pIC50 for each receptor snapshot. Receptor 3 carries the highest mean CNN affinity and pose score. Per-snapshot rank correlation against experiment remains near zero for all conformations.

Ten representative ligands spanning the experimental potency range were evaluated. Both MMGBSA and MMPBSA were computed with full energy decomposition. All decomposed energy components evaluated against Experimental ΔG from the Schrödinger BACE set. MMGBSA VDW achieved the strongest affinity correlation for any component with r = 0.855, *R*^2^ = 0.732, *ρ* = 0.782, and p = 0.002. Benjamini-Hochberg false discovery rate applied reveal MMGBSA VDW as the only statistically significant energy component after correction (adjusted p = 0.010) (Table 2). The VDW range across all 10 compounds ranged from -33 to -52 kcal/mol (Figure 4A). MMGBSA EGB was evaluated with *ρ* = -0.709, p = 0.022, ranging from 65-105 kcal/mol, showing significant inverse rank order correlation. MMGBSA EEL was evaluated with r = 0.393, *R*^2^ = 0.155, *ρ* = 0.382. MMGBSA TOTAL was evaluated with r = 0.529, *R*^2^ = 0.280, *ρ* = 0.382, p = 0.116 (Figure 4A). MMGBSA TOTAL ranged from -10 to -29 kcal/mol vs experimental -7.9 to -11.4 kcal/mol, indicating a scale mismatch between MMGBSA TOTAL and experimental ΔG. MMPBSA showed no correlation against experimental ΔG with r = -0.001 (Figure 4A). It is also worth noting that CAT-4b was evaluated with a very negative MMGBSA TOTAL (-29 kcal/mol) but only moderate potency (ΔG = -9.6 kcal/mol) (Figure 4C).

**Figure 4:**
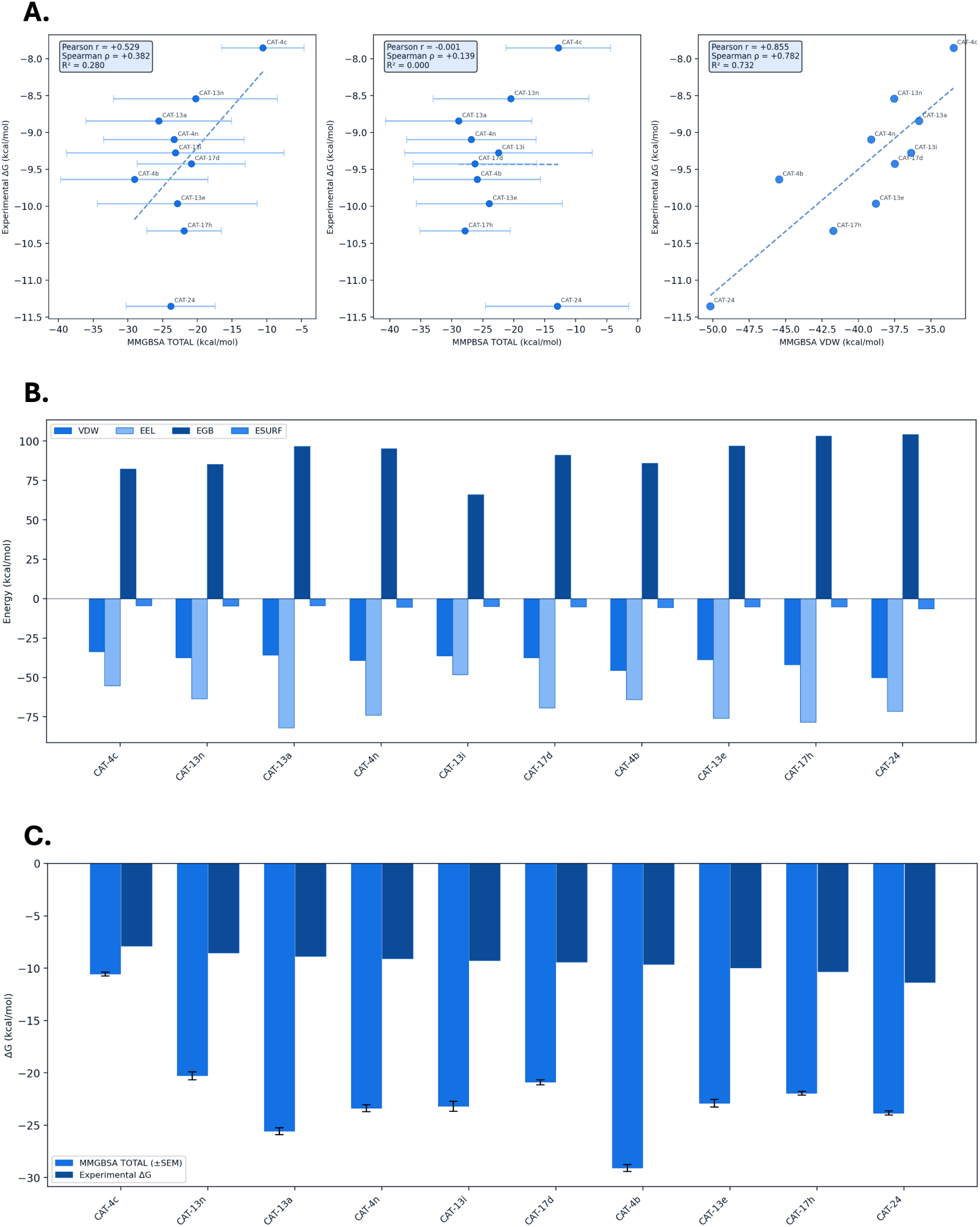
MD/MMPBSA. **A)** MMPBSA and MMGBSA binding energy versus experimental DeltaG (N = 10). Left: MMGBSA Total (kcal/mol) vs Experimental DeltaG (kcal/mol). Middle: MMPBSA Total (kcal/mol) vs Experimental DeltaG (kcal/mol). Right: MMGBSA VDW (kcal/mol) vs Experimental Delta G (kcal/mol). The MMGBSA VDW (right) is the strongest pre-FEP affinity signal (r = 0.855), while total energies are weakened or cancelled by solvation over-correction. **B)** MMGBSA energy component decomposition per ligand (N = 10). Ligands sorted from weakest to strongest experimental binder. **C)** MMGBSA TOTAL vs Experiemtnal DeltaG per ligand. Sorted weakest to strongest experimental binder. The favorable VDW contribution is partially offset by the EGB solvation penalty across the series.

**Table 2:** Benjamini-Hochberg multiple comparison correction for MMPBSA/MMGBSA energy components against experimental ΔG (N=10 ligands).

| Energy Component | Original $p$ | Corrected $p$ | Significant |
| --- | --- | --- | --- |
| MMGBSA VDW | 0.002 | 0.010 | Yes |
| MMGBSA EGB | 0.089 | 0.193 | No |
| MMGBSA EEL | 0.261 | 0.326 | No |
| MMGBSA Total | 0.116 | 0.193 | No |
| MMPBSA Total | 0.997 | 0.997 | No |

A 35-edge perturbation network was constructed across all 36 ligands. A total of 70 simulation legs were evaluated (35 complex legs + 35 solvent legs). All 35 edges were retained for analysis. CAT-13a, with an experimental ΔG = -8.843, was utilized as the anchor ligand in the perturbation network. The network averaged a connectivity of 1.9 edges per ligand. Correlation between raw pairwise ΔΔG values across all 35 edges was then evaluated. Pairwise ΔΔG correlation was observed with r = 0.563, *R*^2^ = 0.317, *ρ* = 0.481, and a regression slope of 1.05. A MAE of 0.98 and an RMSE of 1.27 kcal/mol were observed (Figure 5A). Absolute ΔG calibration via MLE across all 36 ligands was then performed with a correlation of r = 0.662, *R*^2^ = 0.438, and *ρ* = 0.624 obtained. A MAE of 1.02, an RMSE of 1.29, and a mean signed error of -0.228 kcal/mol were observed (Figure 5B). 32 of the 36 ligands were evaluated within 2.0 kcal/mol, with 17 of the 36 evaluated within 1.0 kcal/mol (Figure 5C). A direct comparison of RevFEP was conducted against Schrödinger FEP+ from Wang et al. (2015) on the BACE1 congeneric series. Schrödinger FEP+ reported the following metrics: r = 0.780, MUE = 0.84 kcal/mol, RMSD = 1.03 kcal/mol, and 58 perturbations. RevFEP reported r = 0.662, MAE = 1.02 kcal/mol, RMSE = 1.29 kcal/mol, and 35 edges (Table 4). CAT-24 was observed with the largest difference between experimental and predicted ΔG, with an error difference of 3.2 kcal/mol. In addition, CAT-13b and CAT-4l were observed with errors over 2.5 kcal/mol. Both ligands are single-edged compounds in the perturbation network, with no redundant transformations connecting them to the rest of the network. Uncertainty calibration was then performed, comparing predicted uncertainty against absolute error. The calibration showed little to no correlation with no significance with r = 0.021 and p = 0.903 (Figure 5C). CAT-24 was assessed with *σ* = 0.08 kcal/mol, the lowest uncertainty in the dataset, with an absolute error of 3.2 kcal/mol (Figure 5C).

**Figure 5:**
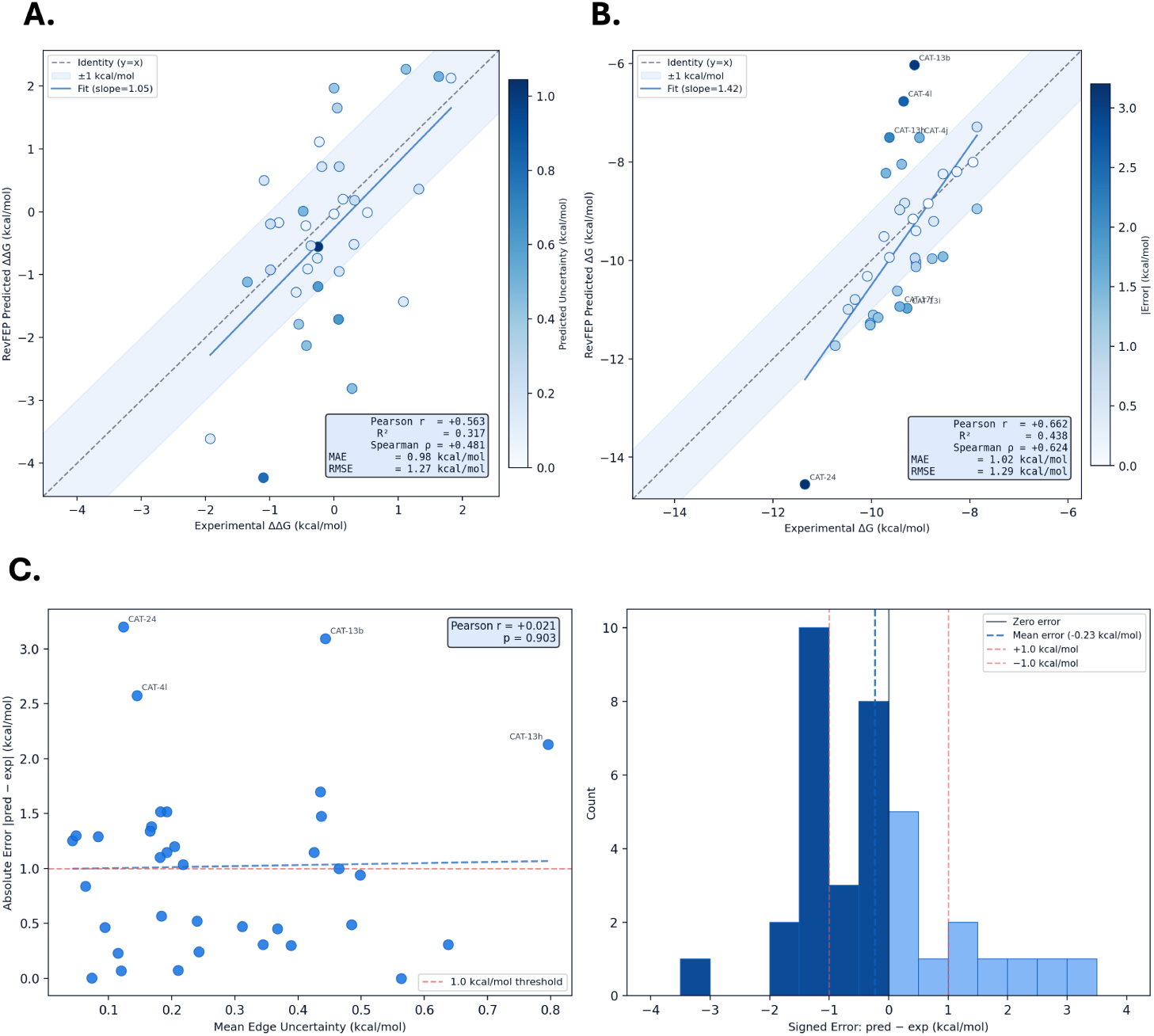
RevFEP calculated using Relative Biding Free Energy. **A)** RevFEP pairwise Delta DeltaG predicted versus experimental for BACE1 (N = 35 edges, raw edge predictions, no MLE). Points are colored by predicted uncertainty. The shaded band indicates ±1 kcal/mol and the fitted slope is 1.05. **B)** RevFEP predicted vs experimental Delta G calibration for BACE1 (N = 36). Calibration curve with fitted slope 1.42. **C)** RevFEP uncertainty calibration analysis (N = 36). Left: predicted uncertainty vs absolute error shows no meaningful correlation (r=0.021, p = 0.903). Right: signed error distribution, with 17 of 36 ligands within ±1.0 kcal/mol.

**Table 3:** RevFEP absolute Δ*G* calibration, MLE network (N=36 ligands).

| Metric | Value |
| --- | --- |
| Pearson $r$ | +0.662 |
| $R^2$ | 0.438 |
| Spearman $\rho$ | +0.624 |
| MAE | 1.02 kcal/mol |
| RMSE | 1.29 kcal/mol |
| Mean signed error | -0.228 kcal/mol |
| Within $\pm 1.0$ kcal/mol | 17/36 (47%) |
| Within $\pm 2.0$ kcal/mol | 32/36 (89%) |

**Table 4:** Direct comparison of RevFEP and Schrödinger FEP+ Wang et al. (2015) on the identical 36-compound BACE1 congeneric series.

| **Metric** | **RevFEP (this work)** | **Schrödinger FEP+** |
| --- | --- | --- |
| N (ligands) | 36 | 36 |
| Perturbations | 35 | 58 |
| Pearson $r$ | +0.662 | +0.780 |
| MAE (kcal/mol) | 1.02 | 0.84 |
| RMSE (kcal/mol) | 1.29 | 1.03 |

**Table 5:** RevFEP uncertainty calibration (N=36 ligands).

| **Metric** | **Value** |
| --- | --- |
| Pearson $r$ (uncertainty versus error) | +0.021 |
| $p$-value | 0.903 |

## 4 Discussion

This study evaluates five computational engines applied to the BACE1 congeneric series, where each engine is benchmarked against the experimental binding free energies to characterize the resolution limits of each engine as well as provide insight into what each engine contributes. The Schrödinger BACE set evaluated in this case study represents a challenging benchmark, requiring computational methods to discriminate potency differences within a tight 3.5 kcal/mol experimental window. The five engines assessed in this study each contribute non-redundant information that informs compound prioritization decisions at successive levels of computational rigor. The following discussion addresses the central question of this study: What does each computational engine contribute individually to the discrimination of a tight congeneric series?

Flexible performed well on pose reconstruction analysis with 97.2 percent of all compounds in the set achieving an RMSD < 2.0 Å and a mean RMSD of 1.16 Å. These metrics validate flexible docking as a reliable engine for pose generation. The exceptional performance on pose reconstruction can be attributed to the open conformation state of the apo protein structure, rendering the active site accessible and favorable for docking Ghosh and Osswald (2014). Affinity correlation performed poorly with a negative physics score (*ρ* = -0.434). This can be explained by the steric bulk from the S1/S3 substituents introducing additional heavy-atom interactions, which drives Vina-style scores down independently of potency Luo et al. (2017). CAT-24 was identified as the only outlier in the pose reconstruction analysis, obtaining an RMSD of 2.36 Å. The poor performance of CAT-24 in Flexible Docking is the first failure observed, as CAT-24 proved to perform poorly across all engine analyses discussed subsequently. This issue suggests that the poor performance of CAT-24 is unrelated to the receptor itself, but rather associated with the geometry of the compound instead. The overall performance of Flexible Docking across this congeneric series suggests that the primary use case of flexible docking is for preliminary pose generation for subsequent downstream analysis. Affinity correlation suggests that this engine is not a reliable source of potency ranking for BACE1 at this resolution.

Boltz CoFolding was identified as the only engine in the pose generation tier to produce a positive correlation direction (r = 0.136, *R*^2^ = 0.018). Taking into consideration the full calibration analysis of Boltz CoFolding, the engine performed poorly overall, with an inability to correctly rank order compounds effectively, and the engine only accounting for 1.8 percent of the variation. The rank ordering limitation may be a result of the tight 3.5 kcal/mol window analyzed. Structural validation confirms that Boltz2 is capable of accurately predicting the receptor conformation in comparison to the co-crystal structure with a protein backbone RMSD of 0.293 Å. The ligand pose RMSD of 4.082 Å significantly exceeds the 2.0 Å threshold indicating that the ligand has been incorrectly placed in the binding pocket. This means that the poor affinity ranking observed is a result of incorrect ligand pose prediction rather than receptor conformation error. It is worth noting that this structural analysis was performed on a single representative ligand on the set, where a full series validation becomes necessary to understand the full picture of accurate pose prediction. When comparing the predicted vs. experimental pIC50 range, a compression in the predicted values has been observed, where more potent molecules have been underpredicted. This is consistent with a known phenomenon of structure-based affinity models on tight congeneric series, where fine potency differences are smaller than typical model error Yang (2026). Regardless of the poor performance of this engine, Boltz CoFolding can be utilized for preliminary pose generation and hedged against flexible docking to determine optimal pose geometry. Calibration of both Flexible Docking and Boltz CoFolding against known inhibitors of the selected target may be the most optimal strategy for selecting which engine to prioritize for pose generation.

Ensemble docking showed Receptor 3 as the best performing receptor conformation. It is worth noting that no receptor snapshot achieved meaningful affinity rank correlation. Receptor performance in the context of Ensemble Docking refers to the pocket accessibility and pose quality from a geometric point of view for downstream FEP setup, rather than affinity rank correlation. A Mean CNN Pose Score of 0.76 suggests that the MD snapshot takes on an open conformation state with ideal conditions for ligand docking Shimizu et al. (2008). Receptor 2 was observed to have the lowest CNN Affinity, indicative of a closed conformation, where the active site is inaccessible for ligand binding. In comparison to Flexible Docking, Ensemble Docking failed to show any improvement in affinity calibration. These results suggest that receptor conformation was not the limitation in flexible docking, rather CNN scoring may be the primary limitation. Taking into consideration both receptor conformation analysis and affinity based analysis, the primary use case of Ensemble Docking is for protein conformation identification, where the goal is to discover which conformation is optimal for docking in a dynamic state.

MD analysis shows MMGBSA VDW as the primary potency driver with the strongest affinity signal in the campaign. This can mechanistically be explained by the shared amidine-containing scaffold facilitating electrostatic and hydrogen bonding interaction with ASP32 and ASP228, leaving substituent variation at the S1/S3 subpockets as the primary source of potency differences Ghosh and Osswald (2014); Genheden and Ryde (2015). Molecular weight vs experimental ΔG correlation yielded a result of r = 0.261, p = 0.466, showing weak correlation and no statistical significance. This confirms that molecular weight is not a meaningful predictor of potency in this subset, and therefore supports the mechanistic interpretation established above. Benjamini-Hochberg correction confirmed MMGBSA VDW as the only statistically significant predictor after correction (adjusted p = 0.010), reinforcing our observation that MMGBSA VDW is the primary potency driver. MMGBSA TOTAL was observed as weaker than VDW alone, largely attributed to the EGB solvation penalty (*ρ* = -0.709, p = 0.022) canceling out the VDW signal. A scale mismatch between MMGBSA TOTAL and experimental ΔG was observed and expected as MMGBSA methods are calibrated for rank ordering rather than absolute ΔG prediction Hou et al. (2011). MMPBSA TOTAL performed poorly overall, where the failure is largely a result of the compounding errors inherent to the method’s numerous calculations and assumptions when applying to ligands with a narrow experimental binding energy range Pearlman (2005). MD with MMPBSA and MMGBSA postprocessing was the first engine in the study to deliver statistically significant affinity signals, as well as marking the transition from geometric to thermodynamic characterization. The equidistant sampling strategy of the 10-compound subset may artificially inflate the variance of the MMPBSA/GBSA metrics, potentially biasing the correlation coefficient upward compared to if performed on a randomly drawn sample. Therefore the reported MMGBSA VDW correlation (r = 0.855) should be interpreted as an upper bound estimate of predicted performance rather than an absolute prediction of the full set. The identification of VDW complementarity with the S1/S3 subpockets as the primary potency driver serves as the guiding pharmacophoric design principle for subsequent DMTA cycle iterations.

RBFE showed the highest affinity correlation of any method across the full 36-ligand series. Pairwise ΔΔG results (MAE = 0.98 kcal/mol) fall at the excellent-to-good boundary established by industry standards, with a regression slope of 1.05 indicating no bias at the edge level. A calibration slope of 1.42 comparing predicted ΔG vs experimental ΔG shows a slight overprediction of potency spread. This is a common characteristic of RBFE as highlighted in Deflorian et al. (2020), where it was established that relative free energy predictions are sensitive to statistical artifacts, especially when experimental bioactivity spans a compressed window. Uncertainty miscalibration shows MBAR *σ* reflects simulation convergence only, as it does not capture force field error, binding mode error, or network topology effects. Network density has been identified as the key improvement layer. The network averaged 1.9 edges per ligand, indicative of moderate connectivity and sufficient for MLE convergence. The MST topology used to construct the network lacks the closed thermodynamic cycles, reducing the ability to detect and correct for systematic simulation errors. In direct comparison against Schrödinger FEP+ on the BACE1 congeneric set, RevFEP achieved competitive performance but fell short with Pearson r = 0.662 and MAE = 1.02 kcal.mol compared to r = 0.780 and MUE = 0.84 kcal.mol for Schrödinger FEP+ Wang et al. (2015). Network density may be a primary explanation for this gap as 58 perturbations were performed compared to 35 edges in this work. The absence of closed thermodynamic cycles as identified above further limits cycle closure corrections which are standard in FEP+ protocols. Nonetheless, RevFEP delivers competitive RBFE performance against industry standards established by FEP+. CAT-13b and CAT-4l were identified as single-edge ligands that both showed the largest errors after CAT-24. As a result, the absolute ΔG estimates for both ligands are dependent on the accuracy of a single pairwise distribution. Additional redundant edges would improve accuracy reliability as there are multiple pathways to iterate over. FEP operates as the terminal layer of this study, where potency rank ordering can be conducted at the highest resolution.

## 5 Conclusion

The comprehensive computational benchmarking study demonstrated here shows that no single engine was able to achieve a sufficient resolution to rank ligands within the 3.5 kcal/mol congeneric window. This benchmarking study evaluates 5 computational engines each contributing key information from pose geometry prediction to thermodynamic characterization, where each engine contributes key decision making factors for synthesis prioritization. Each engine contributes non-redundant information collectively characterizing the resolution limits across different levels of both cost and computational rigor. While validated exclusively on the BACE1 congeneric series, we hypothesize that this resolution hierarchy is likely to generalize to other tight congeneric series as well; however, empirical validation across targets with diverse binding site geometries and ligand chemotypes is required to confirm this. Below is a table summarizing our findings across the different resolution hierarchies.

**Table 6:** Cross-engine affinity correlation summary and direct comparison to Schrödinger FEP+ Wang et al. (2015) on the identical 36-compound BACE1 series.

| Engine | N | Pearson $r$ | $R^2$ | Spearman $\rho$ | MAE (kcal/mol) |
| --- | --- | --- | --- | --- | --- |
| Flexible Docking (CNN) | 36 | -0.155 | — | -0.174 | — |
| Boltz2 Co-Folding | 36 | +0.136 | 0.018 | +0.052 | — |
| Ensemble Docking (CNN) | 36 | -0.229 | — | -0.262 | — |
| MD / MM-GBSA VDW | 10 | +0.855 | 0.732 | +0.782 | — |
| RevFEP (this work) | 36 | +0.662 | 0.438 | +0.624 | 1.02 |
| Schrödinger FEP+ Wang et al. (2015) | 36 | +0.780 | — | — | 0.84 |

With respect to discriminating a large compound set with fine potency differences, this study demonstrates that the computationally inexpensive models evaluated are able to provide structural and conformational insight at lower cost, while thermodynamic engines such as MD and FEP are required for potency discrimination at sub-kcal/mol resolutions.

## Author Contributions

K.A. and C.K. designed the computational campaign and performed the analyses. K.A. and C.C. created the data visualizations and figures. K.A., C.K., and C.C. wrote and revised the manuscript.

## Software and Data Availability

All five engines evaluated are proprietary tools developed by Revilico Inc. All simulation parameters, force field selections, and settings are fully specified in the Methods Section. The Schrödinger BACE1 benchmark dataset is publicly available from Wang et al. (2015). Input files and perturbation network topology are available upon reasonable request to the corresponding author.

## 6 Supplementary Figures

**Figure 6:**
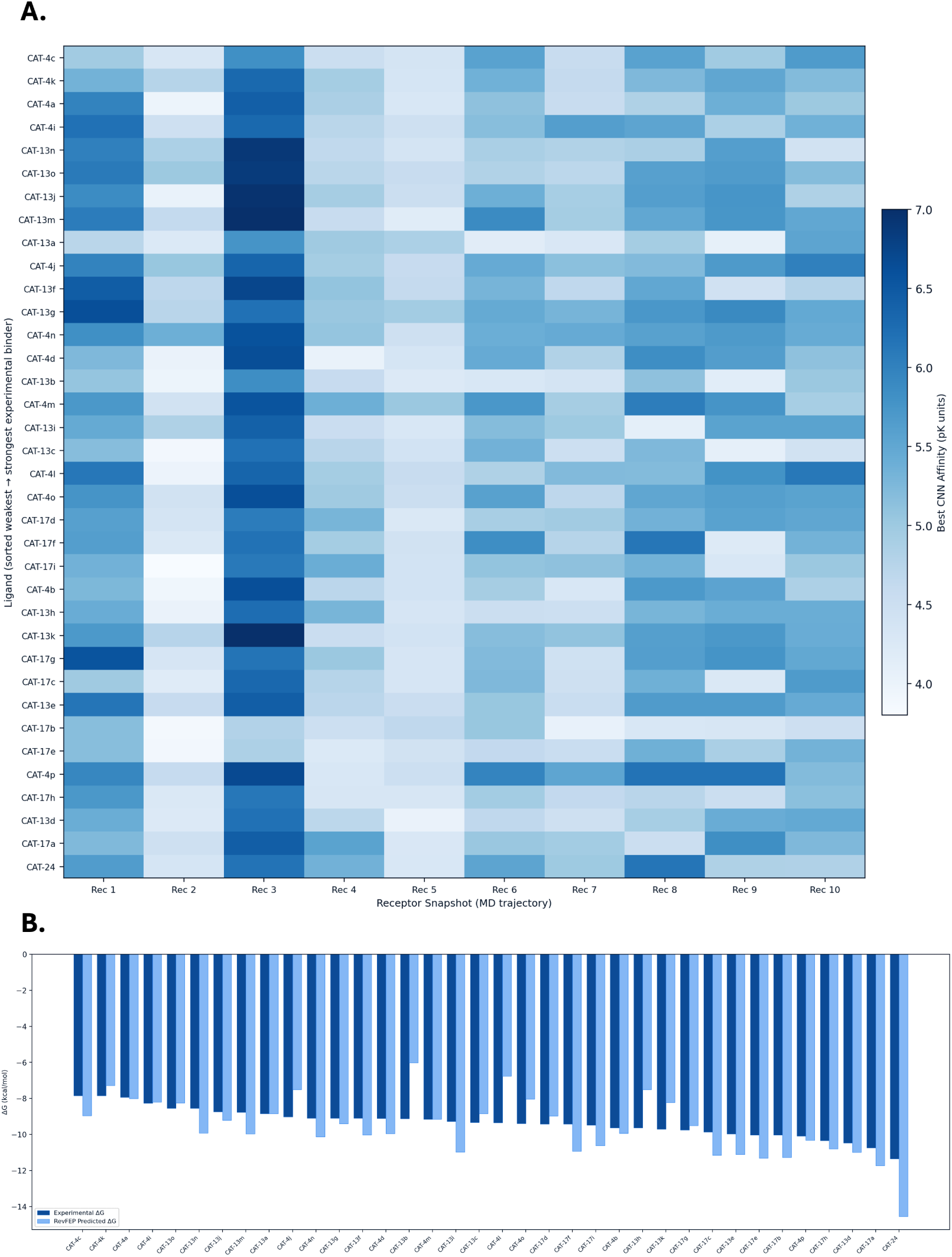
Supplementary Figures. **A)** Ensemble docking CNN affinity heatmap, 36 ligands across 10 receptor snapshots. Receptor 3 shoes the strongest predicted binding across the series. **B)** Per-ligand predicted vs experimental DeltaG sorted from weakest to strongest binder for RevFEP.

## References

Tenchov, R.; Sasso, J. M.; Zhou, Q. A. Alzheimer’s Disease: Exploring the Landscape of Cognitive Decline. ACS Chemical Neuroscience 2024, 15, 3800–3827.

Abdulkhaliq, A. A.; Kim, B.; Almoghrabi, Y. M.; Khan, J.; Ajoolabady, A.; Ren, J.; Bahijri, S.; Tuomilehto, J.; Borai, A.; Pratico, D. Amyloid- and Tau in Alzheimer’s disease: pathogenesis, mechanisms, and interplay. Cell Death & Disease 2026, 17, 21.

Sun, B.; Halabisky, B.; Zhou, Y.; Palop, J. J.; Yu, G.; Mucke, L.; Gan, L. Imbalance between GABAergic and Glutamatergic Transmission Impairs Adult Neurogenesis in an Animal Model of Alzheimer’s Disease. Cell stem cell 2009, 5, 624–633.

Kruse, P.; Eichler, A.; Klukas, L.; Lenz, M. A synapse perspective on the function of the amyloid precursor protein. Science Progress 2025, 108, 00368504251360728.

Zhao, J.; Liu, X.; Xia, W.; Zhang, Y.; Wang, C. Targeting Amyloidogenic Processing of APP in Alzheimer’s Disease. Frontiers in Molecular Neuroscience 2020, 13, 137.

Koelsch, G. BACE1 Function and Inhibition: Implications of Intervention in the Amyloid Pathway of Alzheimer’s Disease Pathology. Molecules : A Journal of Synthetic Chemistry and Natural Product Chemistry 2017, 22, 1723.

Ghosh, A. K.; Osswald, H. L. BACE1 (-Secretase) Inhibitors for the Treatment of Alzheimer’s Disease. Chemical Society reviews 2014, 43, 6765–6813.

Shimizu, H.; Tosaki, A.; Kaneko, K.; Hisano, T.; Sakurai, T.; Nukina, N. Crystal Structure of an Active Form of BACE1, an Enzyme Responsible for Amyloid Protein Production. Molecular and Cellular Biology 2008, 28, 3663–3671.

Patil, R.; Das, S.; Stanley, A.; Yadav, L.; Sudhakar, A.; Varma, A. K. Optimized Hydrophobic Interactions and Hydrogen Bonding at the Target-Ligand Interface Leads the Pathways of Drug-Designing. PLoS ONE 2010, 5, e12029.

Imran, S.; Patel, M.; Noroozifar, M.; Kerman, K. Recent advances towards BACE1 drug discovery and therapeutics design. RSC Medicinal Chemistry 17, 2306–2325.

Wang, L. et al. Accurate and Reliable Prediction of Relative Ligand Binding Potency in Prospective Drug Discovery by Way of a Modern Free-Energy Calculation Protocol and Force Field. Journal of the American Chemical Society 2015, 137, 2695–2703.

Ross, G. A.; Lu, C.; Scarabelli, G.; Albanese, S. K.; Houang, E.; Abel, R.; Harder, E. D.; Wang, L. The maximal and current accuracy of rigorous protein-ligand binding free energy calculations. Communications Chemistry 2023, 6, 222.

Nguyen, N. T.; Nguyen, T. H.; Pham, T. N. H.; Huy, N. T.; Bay, M. V.; Pham, M. Q.; Nam, P. C.; Vu, V. V.; Ngo, S. T. Autodock Vina Adopts More Accurate Binding Poses but Autodock4 Forms Better Binding Affinity. Journal of Chemical Information and Modeling 2020, 60, 204–211.

Passaro, S.; Corso, G.; Wohlwend, J.; Reveiz, M.; Thaler, S.; Somnath, V. R.; Getz, N.; Portnoi, T.; Roy, J.; Stark, H.; Kwabi-Addo, D.; Beaini, D.; Jaakkola, T.; Barzilay, R. Boltz-2: Towards Accurate and Efficient Binding Affinity Prediction. 2025; http://biorxiv.org/lookup/doi/10.1101/2025.06.14.659707.

Liu, Y.; Tang, H.; Niu, T.; Wang, J. A Comparative Study of Deep Learning and Classical Modeling Approaches for Protein-Ligand Binding Pose and Affinity Prediction in Coronavirus Main Proteases. Journal of Chemical Information and Modeling 2026, 66, 731–743.

Totrov, M.; Abagyan, R. Flexible ligand docking to multiple receptor conformations: a practical alternative. Current Opinion in Structural Biology 2008, 18, 178–184.

Huang, S.-Y.; Zou, X. Ensemble docking of multiple protein structures: considering protein structural variations in molecular docking. Proteins 2007, 66, 399–421.

Genheden, S.; Ryde, U. The MM/PBSA and MM/GBSA methods to estimate ligand-binding affinities. Expert Opinion on Drug Discovery 2015, 10, 449–461.

Muegge, I.; Hu, Y. Recent Advances in Alchemical Binding Free Energy Calculations for Drug Discovery. ACS Medicinal Chemistry Letters 2023, 14, 244–250.

Rosignoli, S.; Paiardini, A. Boosting the Full Potential of PyMOL with Structural Biology Plugins. Biomolecules 2022, 12, 1764.

Meli, R.; Morris, G. M.; Biggin, P. C. Scoring Functions for Protein-Ligand Binding Affinity Prediction Using Structure-based Deep Learning: A Review. Frontiers in Bioinformatics 2022, 2, 885983.

Fooshee, D.; Andronico, A.; Baldi, P. ReactionMap: An Efficient Atom-Mapping Algorithm for Chemical Reactions. Journal of Chemical Information and Modeling 2013, 53, 2812–2819.

Lindorff-Larsen, K.; Piana, S.; Palmo, K.; Maragakis, P.; Klepeis, J. L.; Dror, R. O.; Shaw, D. E. Improved side-chain torsion potentials for the Amber ff99SB protein force field. Proteins 2010, 78, 1950–1958.

Mark, P.; Nilsson, L. Structure and Dynamics of the TIP3P, SPC, and SPC/E Water Models at 298 K. The Journal of Physical Chemistry A 2001, *105*, 9954–9960.

Dislich, B.; Lichtenthaler, S. F. The Membrane-Bound Aspartyl Protease BACE1: Molecular and Functional Properties in Alzheimer’s Disease and Beyond. Frontiers in Physiology 2012, 3, 8.

Ries, B.; Alibay, I.; Swenson, D. W. H.; Baumann, H. M.; Henry, M. M.; Eastwood, J. R. B.; Gowers, R. J. Kartograf: A Geometrically Accurate Atom Mapper for Hybrid-Topology Relative Free Energy Calculations. Journal of Chemical Theory and Computation 2024, 20, 1862–1877.

Jakalian, A.; Jack, D. B.; Bayly, C. I. Fast, efficient generation of high-quality atomic charges. AM1-BCC model: II. Parameterization and validation. Journal of Computational Chemistry 2002, *23*, 1623–1641.

Boothroyd, S. et al. Development and Benchmarking of Open Force Field 2.0.0: The Sage Small Molecule Force Field. Journal of Chemical Theory and Computation 2023, *19*, 3251–3275.

Maier, J. A.; Martinez, C.; Kasavajhala, K.; Wickstrom, L.; Hauser, K. E.; Simmerling, C. ff14SB: Improving the Accuracy of Protein Side Chain and Backbone Parameters from ff99SB. Journal of Chemical Theory and Computation 2015, 11, 3696–3713.

Ding, X. Bayesian Multistate Bennett Acceptance Ratio Methods. Journal of Chemical Theory and Computation 2024, 20, 1878–1888.

Luo, Q.; Zhao, L.; Hu, J.; Jin, H.; Liu, Z.; Zhang, L. The scoring bias in reverse docking and the score normalization strategy to improve success rate of target fishing. PLoS ONE 2017, 12, e0171433.

Yang, W. Measuring Affinity Compression. 2026; https://zenodo.org/records/20749570/files/Measuring_Affinity_Compression_Yang_2026.pdf?download=1.

Hou, T.; Wang, J.; Li, Y.; Wang, W. Assessing the Performance of the MM/PBSA and MM/GBSA Methods. 1. The Accuracy of Binding Free Energy Calculations Based on Molecular Dynamics Simulations. Journal of Chemical Information and Modeling 2011, *51*, 69–82.

Pearlman, D. A. Evaluating the Molecular Mechanics PoissonBoltzmann Surface Area Free Energy Method Using a Congeneric Series of Ligands to p38 MAP Kinase. Journal of Medicinal Chemistry 2005, 48, 7796–7807.

Deflorian, F.; Perez-Benito, L.; Lenselink, E. B.; Congreve, M.; Van Vlijmen, H. W. T.; Mason, J. S.; Graaf, C. D.; Tresadern, G. Accurate Prediction of GPCR Ligand Binding Affinity with Free Energy Perturbation. Journal of Chemical Information and Modeling 2020, 60, 5563–5579.

